# Single-cell RNA Profiling Identifies Diverse Cellular Responses to EWSR1-FLI1 Down-regulation in Ewing Sarcoma

**DOI:** 10.1101/750539

**Authors:** Roxane Khoogar, Elizabeth R. Lawlor, Yidong Chen, Myron Ignatius, Katsumi Kitagawa, Tim H.-M. Huang, Peter J. Houghton

**Affiliations:** Department of Molecular Medicine, University of Texas Health Science Center at San Antonio; Greehey Children’s Cancer Research Institute; Department of Pediatrics and Communicable Diseases and Department of Pathology, University of Michigan Medical School; Department of Epidemiology and Biostatistics, University of Texas Health Science Center at San Antonio

## Abstract

Single-cell analyses provide insight into time dependent behaviors in response to dynamic changes of oncogene expression. We developed an unbiased approach to study gene expression variation using a model of cellular dormancy induced via EWSR1-FLI1 down-regulation in Ewing sarcoma (EWS) cells. We propose that variation in the expression of EWSR1-FLI1 over time determines cellular responses. Cell state and functions were assigned using random forest feature selection in combination with machine learning. Notably, three distinct expression profiles were uncovered contributing to Ewing sarcoma cell heterogeneity. Our predictive model identified ∼1% cells in a dormant-like state and ∼2-4% with higher stem-like and neural stem-like features in an exponentially proliferating EWS cell line and EWS xenografts. Following oncogene knockdown, cells re-entering the proliferative cycle have greater stem-like properties, whereas for those remaining quiescent, FAM134B-dependent dormancy provides a survival mechanism. We also show cell cycle heterogeneity related to EWSR1-FLI1 expression as an independent feature driving cancer heterogeneity, and drug resistance.

**SIGNIFICANCE:** We show that time-dependent changes induced by suppression of oncogenic EWSR1-FLI1 induces dormancy, with different subpopulation dynamics, including stem-like characteristics and prolonged dormancy. Cells with these characteristics are identified in exponentially growing cell populations and confer drug resistance, and could potentially contribute to metastasis or late recurrence in patients.

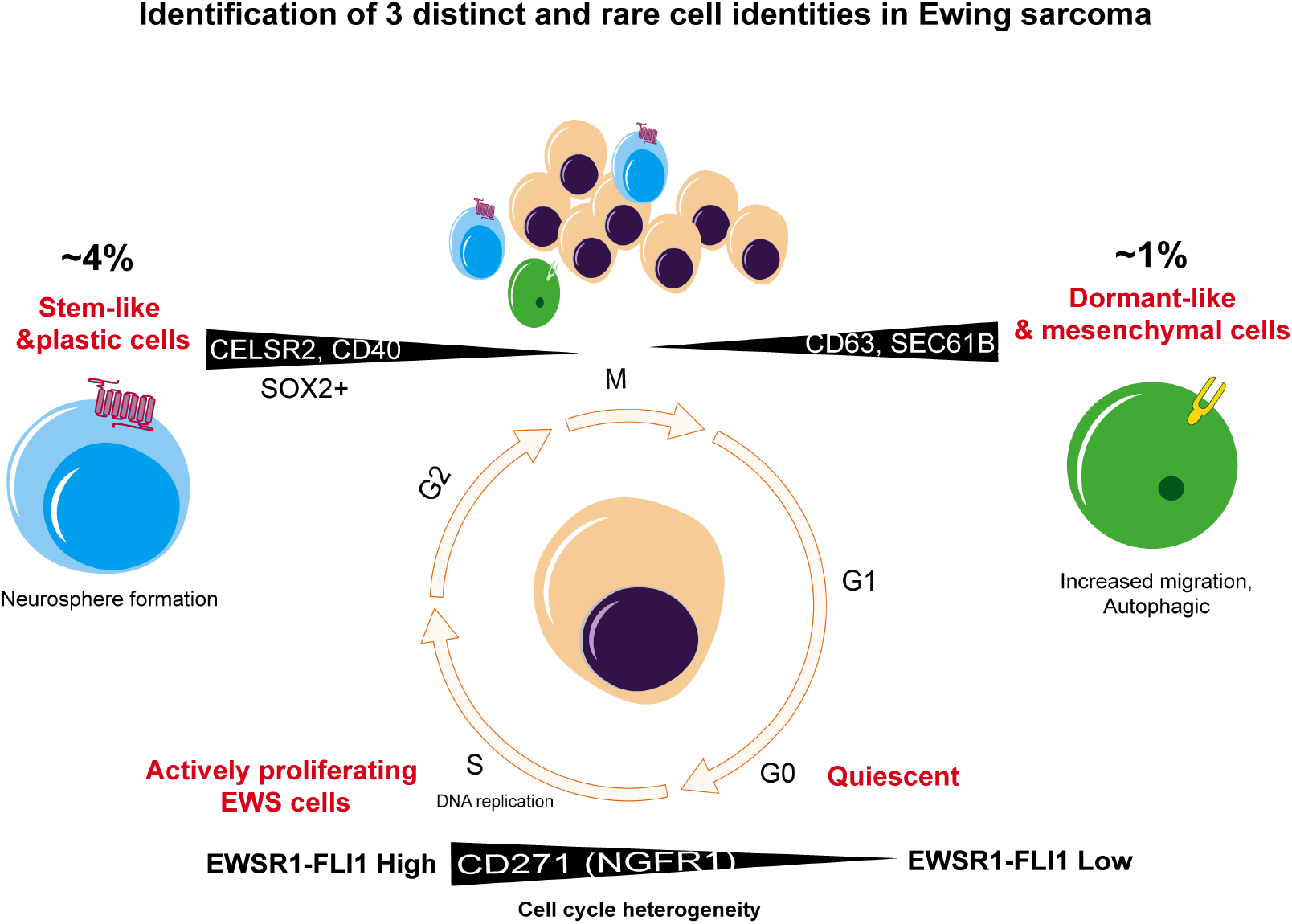

## INTRODUCTION

Stem cell models have been able to accurately describe the ways in which complex hierarchical systems and organs develop, as a result of variation in their basic elements, as in the case of stem cell cycling, illustrated by the capacity for selfrenewal (1), where hematopoietic systems identified the presence of cells with longterm repopulating activity (1–3). Similarly, experiments on spleen formation revealed heterogeneity in the cellular composition of hematopoietic self–renewal and differentiation (1, 2, 4, 5). Environmental factors, such as cytokines, may determine lineage output and multiple differentiation patterns (4–6). For tumors, progression can be described as a complex interplay between induced molecular programs and alterations of gene expression resulting from communication with the environment. Understanding of the diversity of complex molecular programs operational in tumor cells could facilitate the extraction of cellular features and molecular markers that enable better identification of rare subpopulations that contribute to cancer cell heterogeneity and potentially chemotherapy resistance.

The molecular mechanism driving a normal or cancerous cell into a dormant-like or G0 state remains largely undefined. For hematopoietic cells in the bone marrow microenvironment (3) cells may enter a quiescent state or a proliferative state depending on the specific environmental niche (3). The discovery of the micro environmental effect on the fate of stem cells (7–13) gave rise to the concept of “quiescent”-now widely considered to be a “dormant-like cell” in the bone marrow (14, 15). Studies on the interplay between microenvironment and cancer proliferation have suggested additional prognostic factors such as cell cycle and metabolism-related chemo-resistant heterogeneity in cancers (16–18). In T-cell acute lymphoblastic leukemia (T-ALL), for example, various bone marrow niches differentially protect T-ALL cells from chemotherapy by inducing quiescence (18).

In the cancer stem cell model, it is not clear how many sources contribute to the heterogeneity that arises as a consequence of gene expression changes in response to environmental factors (19–22). Thus, these findings underscore the importance of incorporating concepts of regulatory mechanisms that lead to cell plasticity and intra-tumoral heterogeneity, into approaches to stem cell modeling. The advent of singlecell RNA-sequencing technologies has greatly facilitated lineage tracing and sophisticated studies of stem cell heterogeneity when applied to T-cells and stem cells as well as breast cancer cells (23–35). Single cell RNA-seq profiling can reveal gene expression variation between individual cells with important roles in subpopulation dynamics (26–28); for example in the late relapses of patients with Ewing sarcoma (EWS) (29). Many studies have demonstrated heterogeneity within cell populations (30–33), and single-cell RNA-seq technology can provide a promising and unbiased approach to unravel the key molecular programs leading to cancer heterogeneity at cellular resolution (14–18, 34).

Here we have applied single cell sequencing to interrogate heterogeneity in EWS cell populations, with a focus on defining characteristics of cells that may represent a dormant state. Ewing sarcoma is the second most frequent bone cancer in children and young adults, with survival following relapse below 20% (34–38). Despite significant improvements in the success of various therapeutic regimens, relapse remains a critical challenge (39–40). Ewing sarcoma is notable for a significant rate of relapse later than 5-years after diagnosis. For individuals who survived their disease at least 5 years from the time of the diagnosis, the cumulative mortality was 25% at 25 years following entry into the Childhood Cancer Survivor Study. Late relapses in EWS patients may indicate the presence of dormant cells that evade cytotoxic therapy.

In EWS, aberrant transcription of *FLI1* through a reciprocal chromosomal translocation t(11;22) is the most frequent genomic event that places the N-terminus of EWSR1 upstream of the C-terminus of *FLI1*, leading to *EWSR1-FLI1* fusion oncogene. Previous studies have demonstrated that the *EWSR1-FLI1* oncogene can transcriptionally regulate many genes involved in the cell cycle, and those having an effect on survival and differentiation (41–42). Single-cell RNA-seq identifies substantial cell-to-cell heterogeneity for *EWSR1-FLI1* expression and motility gene markers (43). However, it remains largely unknown whether *EWSR1-FLI1* variation can regulate subpopulation dynamics. Thus, it is important to understand how variation occurs in component molecules among seemingly identical cells resulting in diversified population-level responses – ultimately impacting therapy at the single-cell level.

## RESULTS

To experimentally test whether variation in the expression of *EWSR1-FLI1* by individual cells plays a role in subpopulation dynamics, we used an unbiased singlecell transcriptomics strategy. The hypothesis is that high-level expression of *EWSR1-FLI1* will associate with proliferating cells, as it activates expression of E2F (41–42), while low-level expression of *EWSR1-FLI1* will be associated with cellular dormancy, or at least more slowly cycling cells. To generate proliferative quiescence EWSR1-FLI1 was down regulated using RNAi in ES1, ES2, and EW8 Ewing sarcoma cell lines **(Supplemental Fig. S1A)**. To define distinct transcriptional states in response to the stress induced by EWSR1-FLI1 down regulation, first cell proliferation was monitored during 158h following EWSR1-FLI1 knockdown. Down-regulation of EWSR1-FLI1 strongly reduced cell proliferation after 48h, but after a further 120h there was evidence for renewed proliferation in a subset of cells, suggesting a transition from ‘dormancy’ to proliferation **(Supplemental Fig. S1B, C)**. Second, quantitative analysis using a GFP expressing EW8 cell line showed cells transfected with siEWSR1 stop proliferating after 48h **(Supplemental Fig. S1D)**.

To further verify that *EWSR1-FLI1* down-regulation is responsible for the dormant-like state, we knocked down *EWSR1* in *PAX3-FOXO1* fusion positive rhabdomyosarcoma (RH30) cells and we observed no effect on proliferation, whereas it completely inhibited proliferation in EW8 cells (**Supplemental Fig. S1E**). This result suggests that EWSR1-FLI1 has a role in forming the ‘dormant-like’ state. Of note, in EW8 cells knockdown reduced the level of fusion protein, but not the endogenous EWSR1 at 48h, suggesting the fusion protein was more labile. Next we examined cell-cycle distribution in ES1, ES2, EW8, and we observed an increased number of cells in the G0/1 stage following *EWSR1-FLI1* inhibition **(Supplemental Fig. S1F)**. Finally, we used EW8 cells engineered to express EWSR1 that could not be suppressed using the siRNA and we further confirmed that the selective knockdown of EWSR1-FLI1 resulted in a non-proliferative ‘dormant-like’ state (**Supplemental Fig. S1G, H**).

To study changes in gene expression that regulate variable characteristics following *EWSR1-FLI1* down-regulation, the C1 single-cell system (Fluidigm) was used and we performed SMART-seq v4 to prepare libraries from EW8 single cells in three proliferative states (siControl, Dormant 1 and Dormant 2). The first dormant population (D1) was defined as that where cells became proliferatively quiescent (48h time point), and the second population (D2) was obtained at 120h, when a fraction of the cells appeared to recover from the dormant state. Three technically matched cultures with variable *EWSR1-FLI1* expressing population replicates were prepared at three time points (0h, 48h, 120h). An average of 8970 genes were detected in each of 685 viable single cells. Reproducibility was observed between biological replicates (Biological: aggregate R = 0.97416667±0.01) (**Supplemental Fig. S2A-D**), correlations plateauing once ∼ 30 cells had been sampled. In the Dormant 2 (D2; 120h sample), 100 sampled cells reached a similar level of correlation, suggesting greater heterogeneity in this population (**Supplemental Fig. S2E**). The relationship between *EWSR1 or FLI1* with proliferative markers (*PCNA, MCM2, MKI67*) and expression of NGFR (*CD271*) suggest a high correlation between these makers with *EWSR1-FLI1* expression (r= Pearson correlation coefficient) (**Supplemental Fig. S2F**).

### *EWSR1-FLI1* variation and subpopulation dynamics

Transient down-regulation of *EWSR1-FLI1* in EW8 cells led to identification of a bimodal expression pattern for *EWSR1* and *FLI1* genes in the Dormant 2 population. We tracked EWSR1 and FLI1 expression variation in the profiled single-cells obtained from different time points with mean (**μ**), variance (**σ**^2^) and alpha (**α**) parameters. Consistent with the quantitative proliferation studies, a fraction of the cells in D2 revealed lower detected α value than the control **(Supplemental Fig. S3A)**. This suggests that the Dormant 2 population is composed of different subpopulations with different *EWSR1-FLI1* expression.

We considered that individual cells could respond differently to the stress induced by EWSR1-FLI1 down-regulation. To explore the role of EWSR1-FLI1 in coordinating variable responses, principal component analysis (PCA) was performed on the expression values of all detected genes from all cells in three time points **(Supplemental Fig. S3B)**. PCA1 of gene expression profiles from two time points following EWSR1-FLI1 down-regulation distinguished low from high EWSR1-FLI1 expressing cells (Red: D1, 48h from Blue cells: si-Control, 0h). A higher degree of diversity relative to control (Blue cells: siControl, 0h) was observed in the second dormant population (Red cells: D2, 120h). Consistent with the proliferation studies, D2 population (green cells: D2, 120h) was distributed between the two primary populations (Blue cells: si-Control, 0h and Red cells: D1, 48h). This variation may explain the difference in response to *EWSR1-FLI1* contributing to EW8 survival (or ability to repopulate). Exponentially proliferating EW8 cells demonstrated heterogeneous expression of *EWSR1* and *FLI1* (**Supplemental Fig. S3C**), further suggesting the existence of multiple cell states.

To explore the existence of multiple identities within 453 transcriptionally perturbed EW8 cells (Dormant 1+2), a three-step computational pipeline was developed **(Supplemental Fig. S4A)**. The method considered a subset from all detected genes in single dormant cells (1361 genes, 15.2%; P=0.000008) as significant features that coordinate the diverse behaviors in response to EWSR1-FLI1 down-regulation (see methods) (44). Transcriptional states were assigned to each cell allowing two EW8 populations (D1 and D2) to be partitioned into finer sub-populations (**Supplemental Fig. S4B**). Cells were clustered into four distinct subpopulations based on highest silhouette score (**Supplemental Fig. S4C, D**). Interestingly, two predominant clusters (C0 and C2) were identified in the Dormant 2 population **(Supplemental Fig. S4C)**. Subpopulation-level pathway analysis that distinguishes each cluster is presented **(Supplemental Fig. S4D)**.

Next we defined new cell identities by computing an activation score (see methods) for three functional gene panels, cell-cycle regulatory genes, dormancy associated genes and stem cell associated gene features. Following *EWSR1-FLI1* suppression, core stemness genes and dormancy genes showed a strong increase in α at the 120h time point with a significant change in activation score (t test) that characterize each subpopulation **(Fig. 1A, B)**. This allowed identification of different classes of cells with distinct and diverse functions corresponding to *EWSR1-FLI1* fluctuation over time. C0 cells had distinct patterns of expression for genes associated with stemness suggesting re-entry into the proliferative state was associated with a different expression profile being more ‘stem-like’, compared to proliferating cells in the control culture **(Fig. 2A)**. Cluster C0 (“Stem-Like Cluster “) with induced levels of mRNA transcripts for core stem cell genes; Cluster C1 (“Cell-cycle committed Cluster”) with high expression for proliferative markers and low expression for stemlike and survival genes; Cluster C2 (“Dormant-like Cluster”) with high expression for dormancy associated genes; Cluster C3 (Transitional Dormant state”) with undetected proliferative genes, **(Table 1)**. Novel candidate markers from each subpopulation were extracted to uncover driving features leading to heterogeneity in Ewing sarcoma from all single cells (siControl, D1 and D2) **(Fig. 2B)**.

**Figure 1.**
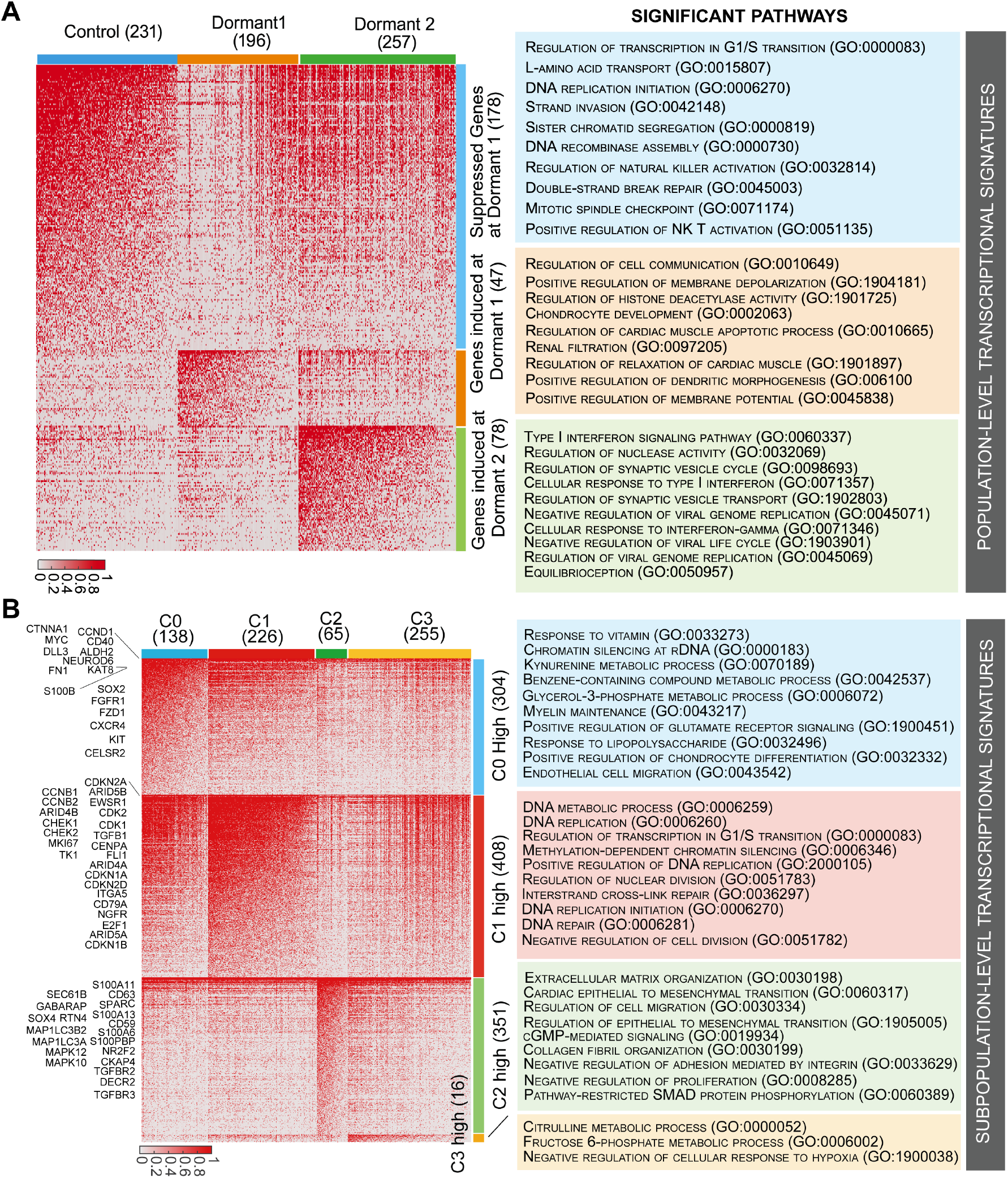
Single-cell RNA-Seq identifies novel transcriptional states when EWSR1-FLI1 is suppressed. Single-cell profiling reveals time-dependent changes in gene expression after EWSR1-FLI1 knockdown. **A**, Population level, genes and pathways suppressed (blue) at 48h (Dormant 1) and 120h (Dormant 2) in EW8 cells, or induced in Dormant 1 (Orange) and Dormant 2 (green) populations (values in parenthesis indicate the number of cells and genes analyzed in each population). **B**, Subpopulation level expression profiles for induced genes in cluster C0 (blue; stem-like), C1 (red; cell cycle committed); C2 (green; Dormant-like), and C3 (orange; transitional dormant). Selection was made using Gene ontology biological process combined score. The score was computed by taking the log of the p value from the Fisher exact test and multiplying the adjusted p-value by the z-score of the deviation from the expected rank (values in parenthesis indicate the number of cells analyzed in each population).

**Figure 2.**
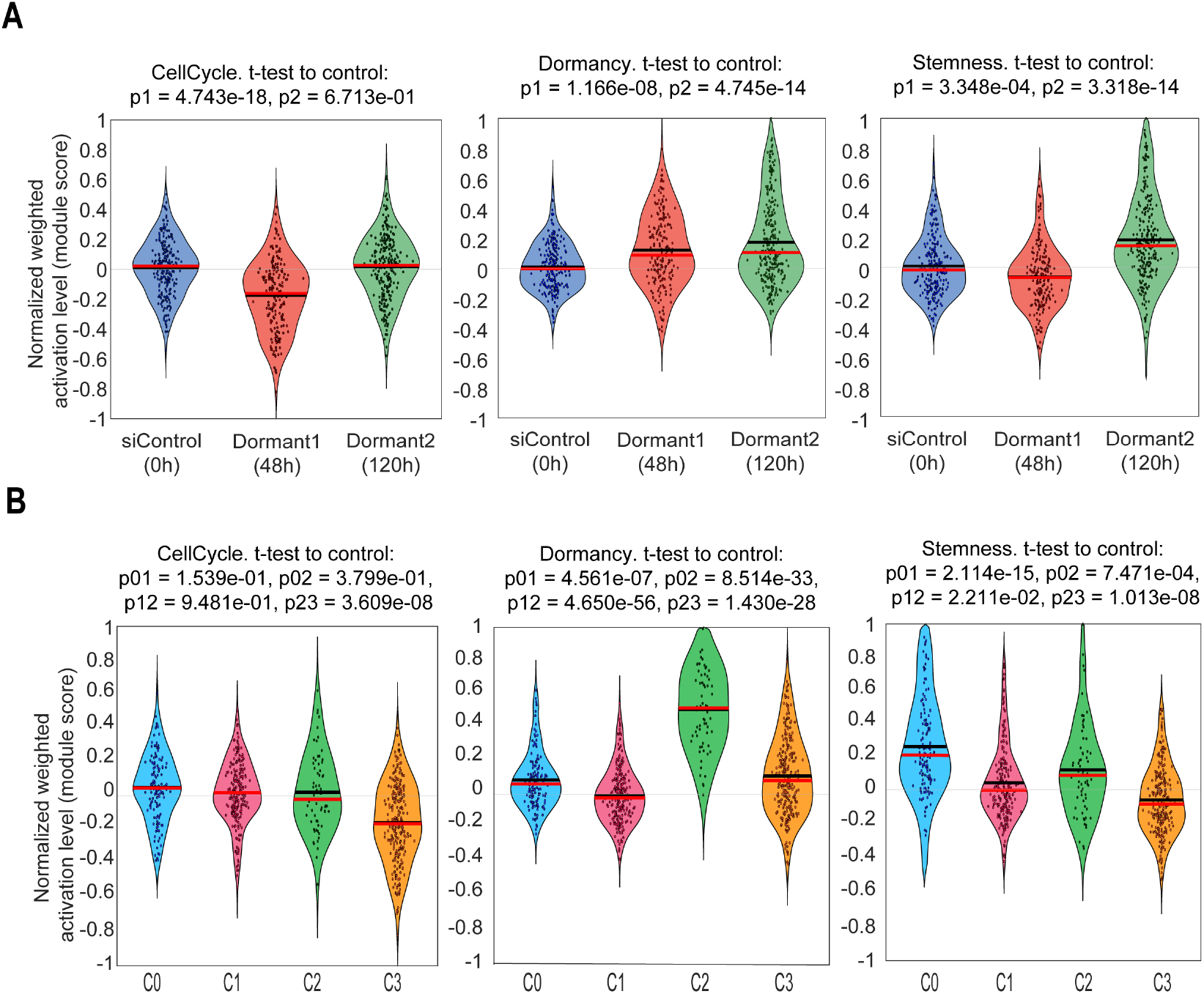
Identification of gene signatures in dormant populations and subclusters of EW8 cells where EWSR1-FLI1 is downregulated. **A**, Gene pathways associated with cell cycle, dormancy and stemness (signatures were obtained from GO) were interrogated in control (siControl) or Dormant 1 and Dormant 2 populations. Genes associated with cell cycle were significantly decreased in Dormant 1 whereas genes associated with ‘stemness’ were increased in Dormant 2 cells. **B**, The same gene sets were interrogated in Dormant 2 cells, and reveal subclusters with distinct characteristics; Cluster C3 shows reduced cell cycle-associated gene expression, whereas cluster C2 demonstrates enhanced gene profile for dormancy, and cluster C0 has elevated levels of ‘stemness’ gene expression.

### Identification of ‘stem-like’ cells in exponentially proliferating EWS cells

To identify stem-like cells in exponentially proliferating cultures of EW8 cells, the single cell mRNA data was searched for cells with the same profile as described for Cluster C0. We classified 232 cells in a heterogeneous population (siControl, 0h) using new class-derived labels. The Neural Network (NN) that obtained 95% predictive precision and 96% success in recall, predicted the transcriptional states in a heterogeneous EW8 cell line (AUC:0.998, CA:0.949, F1:0.954). Four cells from 231 cells (1.73%) in the siControl population demonstrated the stem-like molecular signature (C0) described in **(Fig. 3A)**. Based on genes that identify ‘stem-like’ properties, stem-like cells (C0) were increased from 1% in an unperturbed EW8 population (siControl, 0h) to 58% in the D2 population (siEWSR1, 120h) **(Supplemental Fig. S5A)**. Of note, the C0 cluster showed similar expression levels of *EWSR1* and *FLI1* with the C1 cluster, **(Supplemental Fig. S5B)**. This suggests that *EWSR1-FLI1* fluctuation has led to two distinctive transcriptional states (C0 and C2) after stress induced via *EWSR1-FLI1* suppression and cell-cycle arrest in EW8 cells.

**Figure 3.**
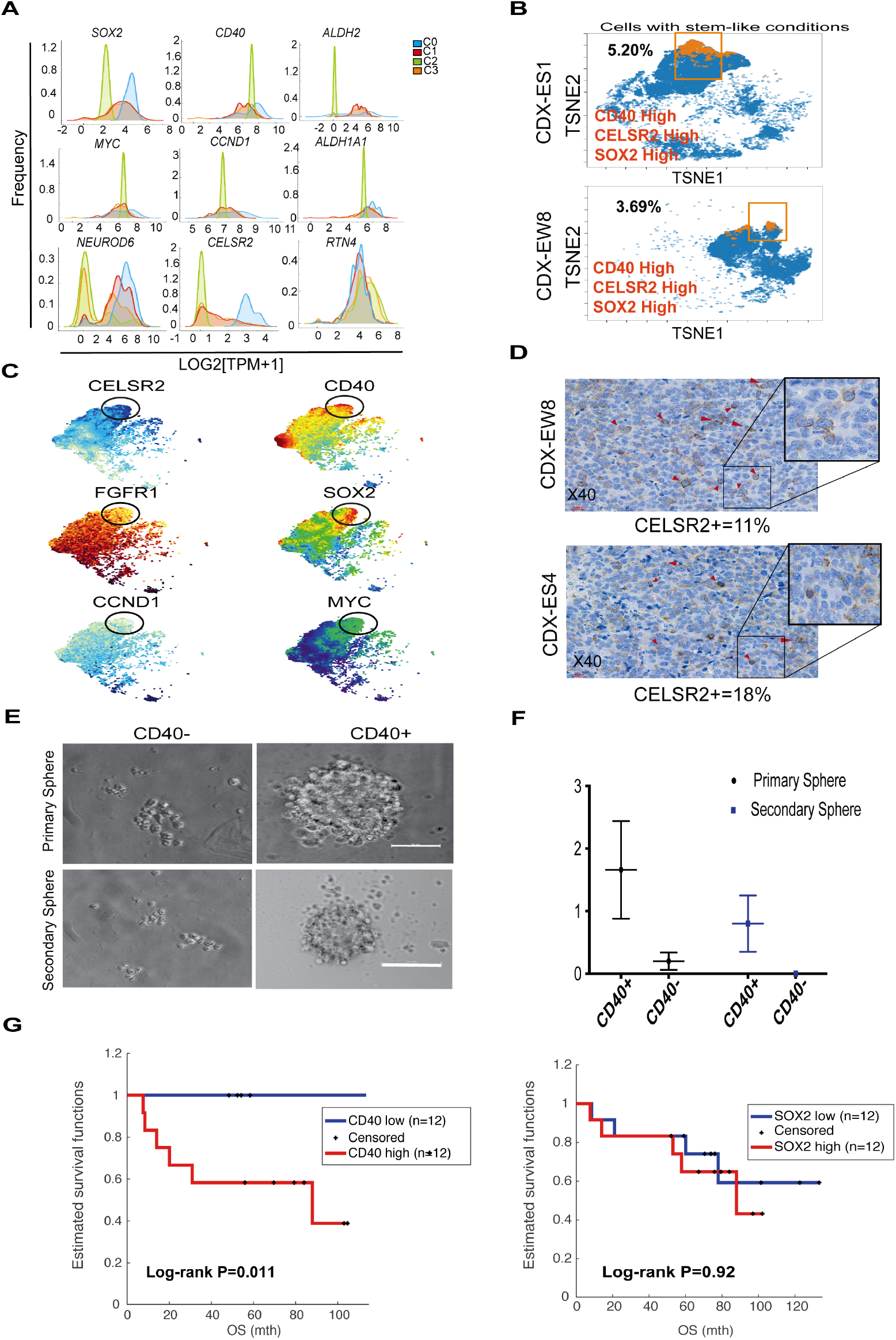
Characterization and validation of distinct subpopulations in Dormant 2. **A**, Discriminative gene-features that define subclusters **B**, (**VISNE PLOTS**) show the low percentage of isolated cells that were considered to express all three proteins at high levels in EWS xenograft tissues. **C, (VISNE PLOTS)** Visualization plots of exponentially proliferating EW8 cells highlight coexpression patterns of newly identified stem-like cell surface markers (CELSR2, CD40) with SOX2, MYC, FGFR1, and CCND1. **D**, Heterogeneous expression of CELSR2 in tissue sections of EWS xenografts. Percentgae of cells scoring positive was assessed from scoring 3-high powered fields (40X). **E**, 7-day self-renewal assayTumor-sphere formation was quantitated and measured from a total of 2400 cells expressing high or low CD40 (Total=4800 cells); (p-value=0.00, unpaired t test). **F**, Quantitation of primary and secondary spheroid colonies formed by CD40-positive and CD-40 negatice EW8 cells. (mean ± SD) **G**, Patient overall survival relative to expression of CD40 and SOX2 ‘stem-like’ markers in Cohort COG GSE63155.

Key signaling pathways characterizing CELSR2/ SOX2/ CD40 positive stem-like cells, were involved in regulating insulin resistance, drug resistance, cytokine-cytokine receptor, PI3K-signaling pathways, *WNT* signaling, and axon guidance **(Supplemental Fig. S6A)**. These findings lead us to profile single-cells based on CD40 expression and further analyze stem cell markers (*ALDH1A1, ALDH2, BRIX1, CCND1, CDC4, CDK1, COL21A1, CTNNA1, CXCR4, DLL3, FGFR1, FN1, FZD1, KAT8, KIT, MYC, NEUROD6, PARD6B, PTEN, SOX2, S100B*). Single cell differentially expressed gene analysis revealed that stemness gene features were more pronounced in the fraction of cells with higher *CD40* levels **(Supplemental Fig. S6B)**.

Among the most highly associated markers, we focused on CD40 and CELSR2, as they appeared to show the most attractive features for a stem like condition. Recovery from EWSR1-FLI1 suppression was associated with induction of inflammatory response genes, immune response genes and paracrine signaling genes in the Dormant 2 population, suggesting a key role for these processes in the fraction of cells with the ability to recover from the Dormant1 state **(Supplemental Fig. S6C)**.

We next sought to determine the frequency of cells co-expressing CD40 and *CELSR2* protein in the unperturbed EW8 cell population and in two cell line-derived EWS xenograft (CDX) models in *scid* mice. Using CyTOF a fraction of cells (∼4%) within EW8 control population (0h), or CDX-ES1 and CDX-EW8 with positive expression for CD40, and CELSR2 was identified and these cells also showed higher levels for SOX2 **(Fig. 3B)**. The mean protein expression levels for CCND1, CD40, CELSR2, CD79a, FGFR1, MYC, SOX2, and KI67 in the rare stem-like subpopulation of EW8 was determined with CyTOF **(Fig. 3C)**. We also examined the cellular heterogeneity for CELSR2 in EWS xenograft tumor sections and identified the expression of CELSR2 only in a small fraction of cells **(Fig. 3D)**. Temporal changes in the expression of a fraction of cells identified with high stemness markers (CD40, CELSR2, and SOX2) at a single cell level with CyTOF revealed CD40 and SOX2 down regulation in the Dormant 1 population was consistent with the transcription profiles and protein expression distribution showing heterogeneity in the expression of all three markers over time using μ and α parameters **(Supplemental Fig. S7A)**.

We aimed to assess the stem-like capabilities of cells with high expression of CD40. From the FACS analysis, we estimated ∼ 5% of EW8 cells were CD40 positive, whereas >90% were CD40 negative. To assess self-renewal capacity, cells FACS sorted for high or low expression of CD40, were plated to form spheroids in neurosphere medium. Formation of spheroids was assessed after 7-10 days. To determine ‘self-renewal’ capacity, a stem-like characteristic, spheroids were disaggregated and cells re-seeded under the same conditions to assess secondary spheroid formation **(Figure 3E)**. The results show that CD40-positive cells formed spheroids more readily than CD40-negative cells, and further, maintained the ability to generate secondary spheroids, indicating self-renewal capabilities **(Figure 3F)**.

### Heterogeneity in ER-autophagy response following EWSR1-FLI1 downregulation and identification of a dormant-like state

Single-cell gene expression data show a distinct expression pattern for marker CD63 in cluster C2 (Dormant-like cluster) (μ_C0_=9.66±0.5; μ_C1_=9.18±0.61; μ_C2_=10.88±0.68; μ_C3_=9.73±0.82, P=0.000, ANOVA) **(Fig. 4A)**. Using the Spearman correlation, the relationship between cell surface marker CD63, a member of the tetraspanin family of membrane proteins, and dormancy associated gene features (TGFBR2, TGFBR3, CDKN2A) was explored **(Fig. 4B)**. Consistent with a previous report (43), we observed higher mean level of motility genes (*FN1, SERPINE2, TIMP1*) and epithelial to mesenchymal regulatory genes (*ITGA4, CALD1, COL3A1, COL5A2, FN1, GSN, MYL6, MYL12A, SPARC, TWIST1, VCL*) in cluster C2 (t-test, P<0.001-0.000001) **(Supplemental Fig. S8A)**, suggesting C2 cells transitioned to a more mesenchymal and motile state following EWSR1-FLI1 down-regulation.

**Figure 4.**
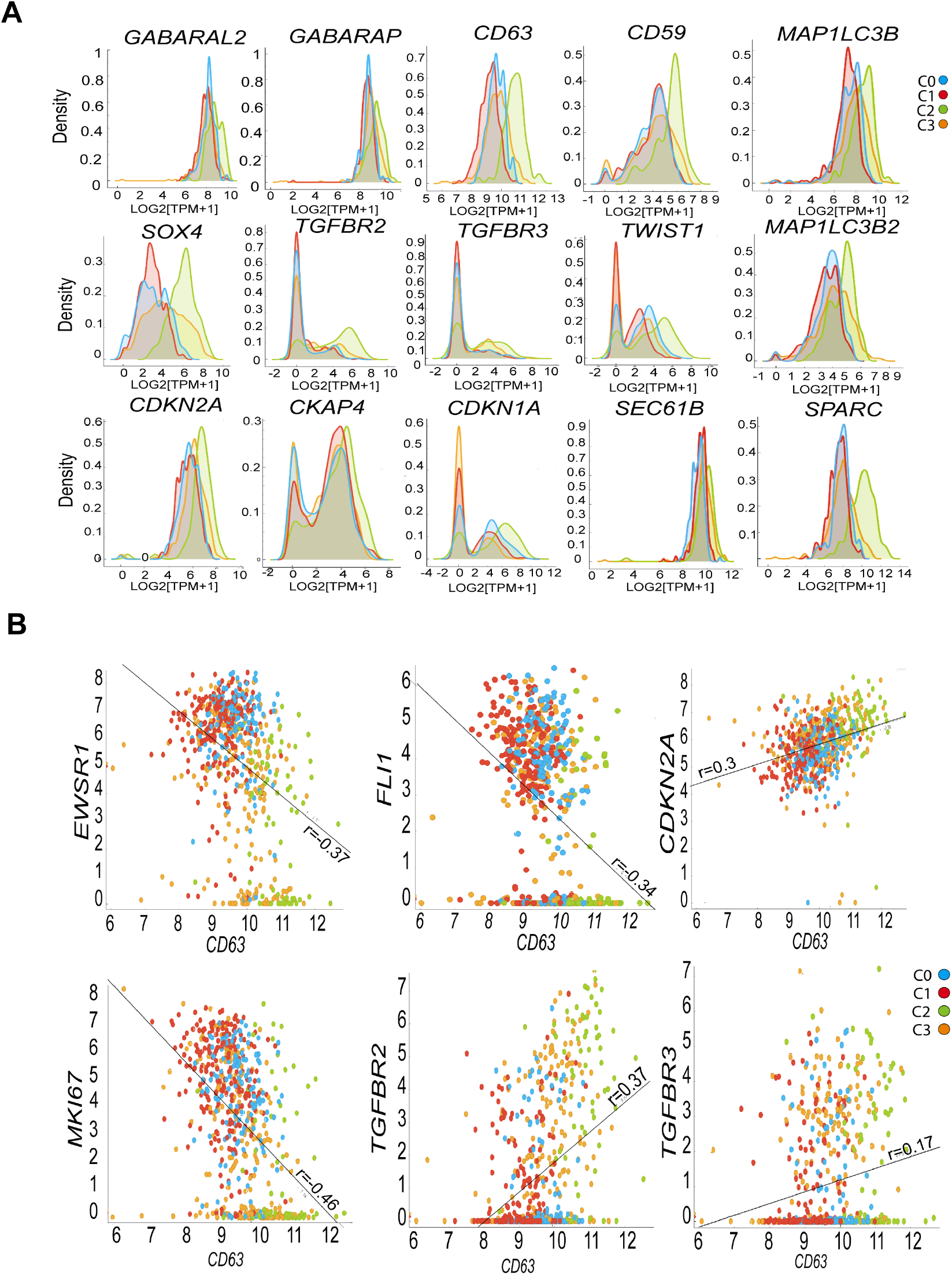
Expression profiles characterizing C2 cluster with Dormant-like features. **A**, C2 in Dormant 2 populations clusters are characterized by increased transcripts for autophagic genes (*GABARAP, MAP1LC3B, MAP1LC3B2*) and TGFβ receptors and cyclin-dependent kinase inhibitory genes (*CDKN1A, CDKN2A*). **B**, The relationship between *CD63* marker and dormancy associated gene markers (*TGFBR2 and TGFBR3*), proliferation markers (MKI67, CDKN2A) and *EWSR1* and *FLI1* in each subpopulation (Pearson correlation). (C0: Stem-like (blue), C1:Committed to the cell-cycle (Red), C2;Dormant-like (Green), C3;Transitional dormant state (Orange).

Population-level protein study of EW8 cells revealed bimodal expression values for dormancy-associated gene markers (*TGFBR2, CDKN2A*) **(Fig. 5A)** following *EWSR1-FLI1* down-regulation. The variation in the expression levels of dormancy associated gene features in the dormant states (48,120h) was validated using CyTOF with previously described parameters. This single-cell protein study confirmed that cells in the proliferative quiescent populations show induced levels of dormancy associated gene features, *TGFBR2* (μ=172.1±27.2;α _D1, D2_=34.91%, 12.92%) and *CDKN2A* (μ=12.83; α _D1, D2_= 37.35%, 11%) in a larger fraction of cells relative to control *TGFBR2* (μ=126.97, α _c_= 2.7%) and *CDKN2A* (μ=11, α _c_= 2.07%) **(Fig. 5A-C)**. T-SNEvisualization plot shows increase in the fraction of cells with dormant conditions (D1, ∼36%), and recovery from a dormant state (D2, ∼12%) when we cluster cells based on their coexpression patterns for both dormancy-associated markers in each population (C, D1, D2) **(Fig. 5C)**. Using CyTOF we next explored co-expression of TGFBR2, CDKN2A with CD63 in cells isolated from two EWS-cell line derived xenografts (CDX-ES4, CDX-EW8). Approximately 1% of isolated cells were detected with positive expression for all three features **(Fig. 5D)**. These data are consistent with the class-derived label prediction and suggests that CD63 can mark rare cells with dormant features. We noted a distinct expression pattern for the growth arrest-specific family members (*GAS1, GAS2L1, GAS5, GAS2Ls*) in C2 cells, whereas *DECR2, MAPK10* (JNK3A), MAPK12, *TWIST, NR2F2*, and survival genes (*S100A11, S100A1S, S100A6, S100A10, S100PBP*) levels were increased in C2 (dormant-like subpopulation), further underscoring the altered transcriptional state in C2 **(Supplemental Fig. S9A)**. Consistent with our dormant model induced via *EWSR1-FLI1* down-regulation, single cells identified based on their relative transcript number for CD63 showed statistically significant deferentially expressed genes involved in the cell-cycle, senescence and drug resistance pathways **(Supplemental Fig. S9B)**.

**Figure 5.**
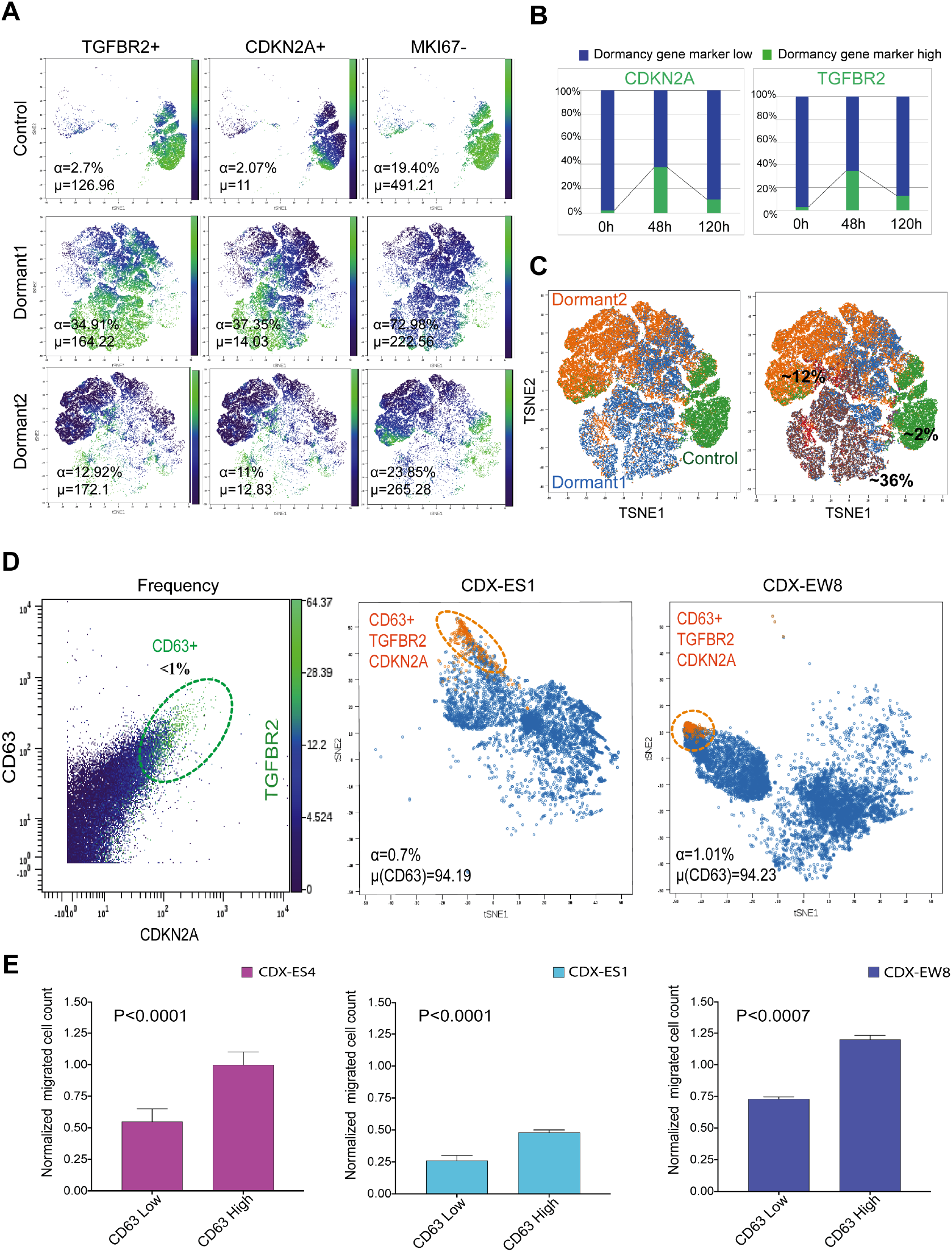
Identification and characterization of cells with dormancy characteristics in cell cultures and xenografts. **A**, Enrichment of a fraction of EW8 cells with positive dormancy gene features (*CDKN2A, TGFBR2*) determined by CyTOF in the Dormant 1 and Dormant 2 populations, showing bimodal kinetics over time. **B, (VISNE PLOTS)** overview of the shift in the frequency of the positive dormant cells over time (120h). **C**, Distribution of cells with dormant features (TGFBR2+, CDKN2A+, MK167−) in siControl, Dormant 1 and Dormant 2 populations combined. **D**, *Left*, Illustration of the gating strategy for detecting cells with high levels of the membrane marker CD63 that are also expressing dormant features (high TGFRB2 and high CDKN2A). CD63-positive, TGFRB2^High^, CDKN2A^High^ represents <1% of the population of exponentially growing EW8 cells. *Center, and right panels*, visualization of the rare fraction of cells isolated from dissociated CDX-ES1 and CDX-EW8 xenografts with dormant characteristics with ViSNE plots. E, Cells dissociated from ES4, ES1 and EW8 xenografts were FACs sorted for high or low CD63 expression, and their ability to migrate was determined with the chemotaxis assay. (P-values are from unpaired t-test).

Under the dormant condition induced by *EWSR1-FLI1* down-regulation, a network of genes regulating selective autophagy (*FAM134B, GABARAP, MAPL3CB*) and ER response to stress (*CD63, SEC61B, RTN4, CLIMP-63*) in the sub-cluster C2 was identified as genes differentiating this cluster **(Supplemental Fig. S9C)**. Variation in the component genes regulating autophagy and ER protein expression reflects a discriminative feature for dormant-like state cluster 2. This cluster is defined by a failure to re-enter the cell cycle (at 120h) but with common gene expression activity for survival genes with cluster C0, the stem-like population (*S100BP, S100PP, S100A10, S100A11, S100A13, S100A6, COX6A1, COX6C, COX7A2, COX8A*). To explore migration of EWS cells using a chemotaxis assay, cells were isolated from EWS xenografts and sorted by FACS based on the number of CD63 surface molecules per cell. The results showed that migration was higher in the CD63 positive (t-test, P<0.0001, n=3), **(Fig. 5E)**.

Co-localization of FAM134B with MAPL3CB was strongly increased in the induced dormant model, after 48h of *EWSR1-FLI1* down-regulation (**Fig. 6A, B**). Furthermore, dormant cells also exhibited increased expression of lysosome marker, CD63 compared to control. These observations support the idea that cells with lower *EWSR1-FLI1* levels which remained in a dormant state for an extended period of time showed higher levels of gene features involved in the autophagy process, as CD63 has been shown to associate with FAM134B and MAP1L3CB in autophagic cells (45). The role of EWSR1-FLI1 in ER activity was investigated by monitoring the levels of the ER-resident protein SEC61B. Increased expression of *SEC61B* was observed in dormant cells (C2) with low levels of *EWSR1-FLI1*. The presence of expanded ER, especially in the cell periphery was further confirmed in dormant cells with lower FAM134B levels by IHC, (**Fig. 6C, D**). These data thus show the link between a dormant model induced via EWSR1-FLI1 suppression and autophagy. Therefore, we concluded that ER-autophagy is a distinguishing cellular behavior contributing to cell survival in a prolonged dormant-like state (C2) in EW8 cells.

**Figure 6.**
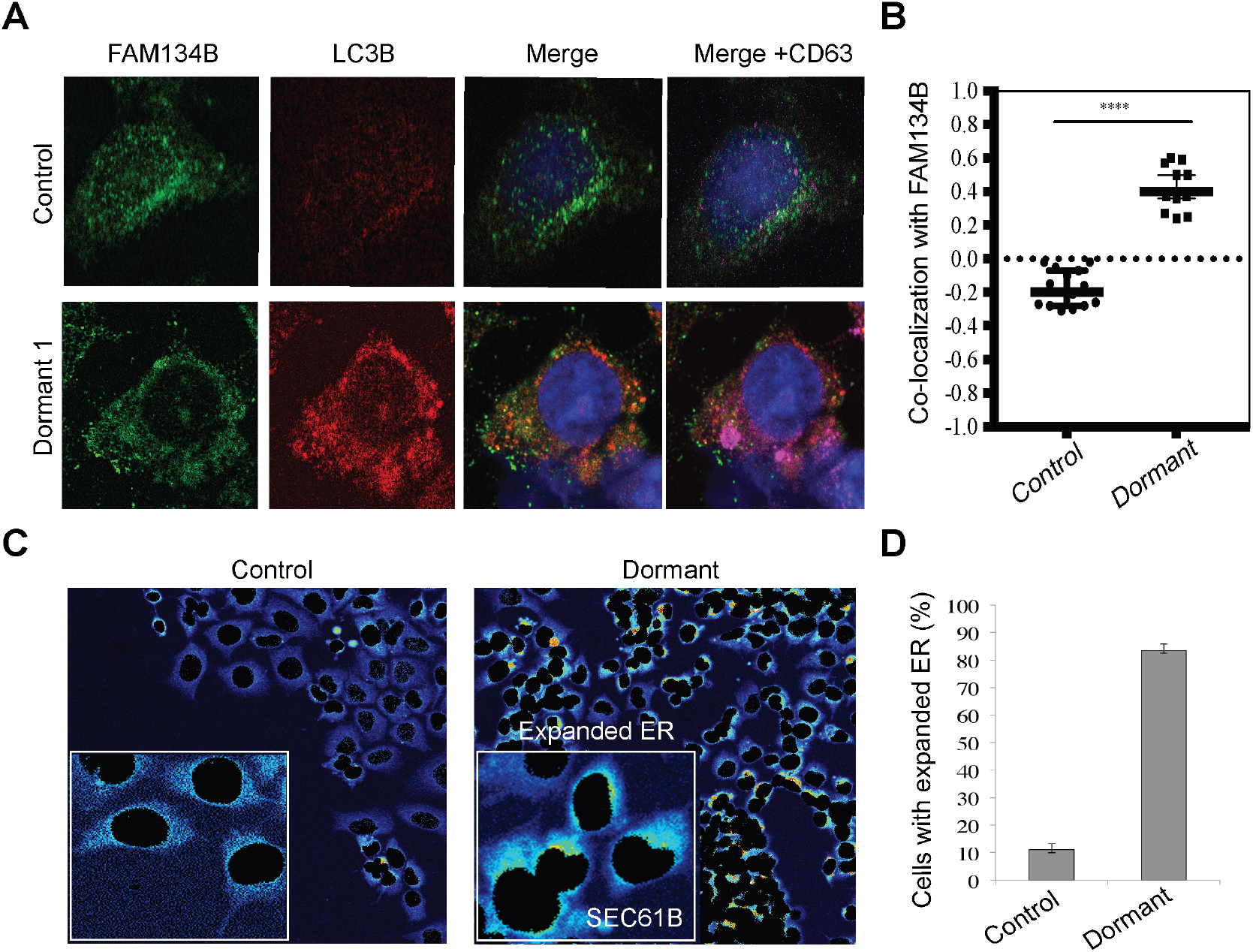
EW8 cells possess altered selective autophagy activity and expanded ER. **A**, Fluorescence images of EW8 cells transfected with siEWSR1 or siControl; EW8 Cells were fixed after 48h immunostained and subsequently cells were imaged via fluorescent microscopy to study the induction of the autophagy marker LC3B and lysosomal marker CD63 when EWSR1-FLI1 is suppressed. **B**, Quantification of FAM134B co-localization presented as Pearson’s correlation coefficient (r); ****P < 0.0001, t-test, n=15 fields (∼150 cells) in the dormant cells shown in A using the colocalization tool in imageJ. **C**, EW8 cells were treated with siEWSR1 or siControl, before being stained for ER resident protein SEC61B. The expansion of ER is shown following EWSR1-FLI1 down regulation (boxed areas). **D**, Quantification of cells with expanded ER after masking nuclei. **** P, 0.0001, t-test, error bars indicate s.d., n=150 cells.

### Variability in the expression of Nerve Growth Factor (NGFR-CD271)

To determine whether EW8 cells with either high or low levels of *EWSR1-FLI1* transcripts had different functional characteristics, we identified the nerve growth factor receptor (*NGFR; CD271*) as a marker strongly correlated with the expression of *EWSR1-FLI1* across all experimental samples and time points (μ_Control_=.3±1.86; μ_Dormant1_= 0.8±1.52; μ_Dormant2_=1.53±1.8, r = 0.41, P<0.0000, ANOVA) **(Supplemental Fig. S10A)**. Further experimental validation with CyTOF confirmed that NGFR (CD271) was expressed heterogeneously in EW8 cells and down regulated following EWSR1-FLI1 suppression **(Fig. 7A)**. Similarly, FACs analysis showed decreased surface expression of NGFR 48h after EWSR1-FLI1 knockdown **(Figure 7B**, *left panel*) NGFR is a neuroectodermal stem cell marker (46) and ectopic EWSR1-FLI1 overexpression resulted in a neural phenotype in rhabdomyosarcoma cell line RD (47). Our data revealed that actively proliferating cells show positive correlation between EWSR1-FLI1 expression and CD271 or nerve growth factor receptor 1 (NGFR1), whereas stem-like cells (C0) demonstrated higher levels of NGFR following EWSR1-FLI1 inhibition (48). This may suggest that cell-cycle activity regulates the expression of CD271 (NGFR1) and NGFR1 can be used as a cell surface marker for identification of cells in a quiescent state. To assess the association between *CD271* expression and EW8 cell proliferation, cells were treated with palbociclib, a CDK4/6 inhibitor to block cell cycle in G1-phase. After 24h surface CD271 was determined by FACS. Palbociclib treated cells presented lower levels of surface CD271 (**Fig. 7B**, *right panel)* suggesting that CD271 expression was cell-cycle related, and that cells isolated based on CD271 expression from the untreated EW8 cell population may be in different phases of the cell cycle or have different proliferation kinetics. Cells were isolated by FACS sorting based on the number of CD271 surface molecules. Isolated cells were binned into four profiles representing the level of surface expression of CD271. Validating our mRNA data the number of cells with high expression for NGFR decreased following EWSR1-FLI1 suppression (**Fig. 7C**).

**Figure 7.**
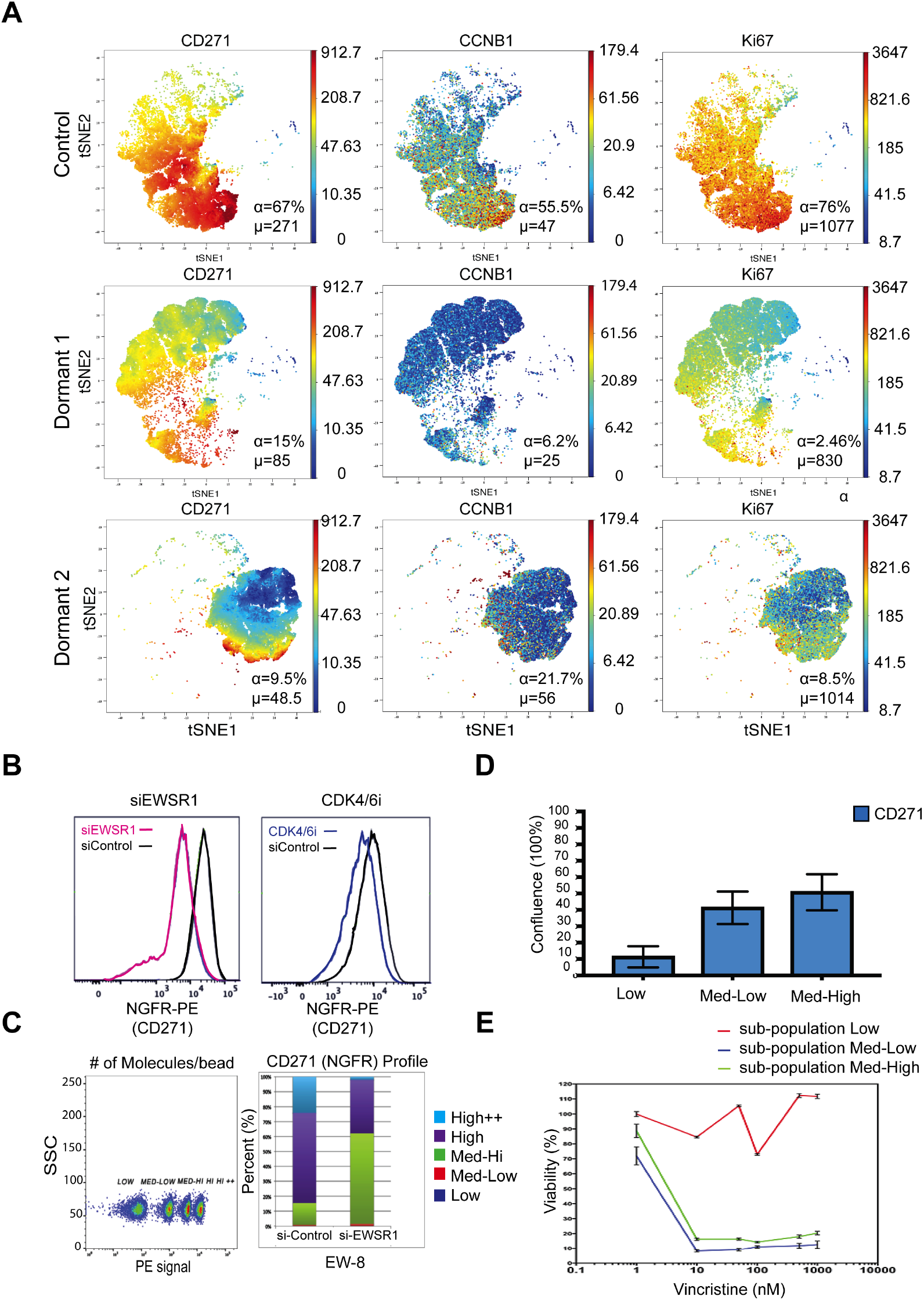
Profiling NGFR expression at a single cell level with CyTOF and identification of a subpopulation with altered sensitivity to vincristine. **A, (VISNE PLOTS)** Visualization of time-dependent behaviors of NGFR (CD271), CCNB1, KI67 expression and variability in the expression of NGFR (CD271-149Sm) when EWSR1-FLI1 is suppressed at two time points, 48h, and 120h (α: The fraction of cells positively expressing the identified marker, μ: the average value of the detected signal for all cells within each window for that marker). **B**, FACS analysis of EW8 cells stained for NGFR (CD271) when cells were transfected with siEWSR1 for 48h (*left panel*, purple line) or treated with CDK4/6 inhibitor (*right panel*, blue line) for 24h **C**, Subpopulations of EW8 cells with different levels of NGFR (CD271) (Population 1= Low; Population 2= Med-Low; population 3= Med-High; population 4= High; Population 5= H++) and quantification of changes in NGFR (CD271) expression in fixed sub-populations based on PE quantibrite beads in response to EWSR1-FLI1 down-regulation. **D**, Quantification of proliferation when EW8 cells were sorted based on CD271 expression. Cell confluence was measured after 96h (Incucyte) and shows the association between proliferative activity and CD271 expression (Sub-population Low= 1.69%; Sub-population Med-Low= 15%; Subpopulation Med-High=26.66%). **E**, EW8 cells with low NGFR (CD271) are resistant to vincristine. Viability was assessed with cyanine deoxyribonucleic acid (DNA) binding dye. The ratio of dead cells at identical time points were normalized to time point 0 value for subpopulations treated with or without vincristine.

To further functionally characterize the CD271 medium-low versus high expression clusters of EW8 cells, cells from each subpopulation were plated with equal densities and allowed to proliferate for 4 days. This proliferation study revealed that the cells with low expression of CD271 did not proliferate, hence remained in a dormant-like state while cells medium-high or high CD271 were highly proliferative. The lack of proliferation in the CD271-low cells, suggests that these cells are proliferatively quiescent **(Fig. 7D)**.

### Cells with Low Expression of CD271 are resistant to vincristine

We next explored the cellular response to vincristine. We hypothesized that cells with low CD271 levels (subcluster-Low) would be resistant to the antimitotic agent vincristine, by virtue of the lack of proliferation. Cells were plated at equal densities, and exposed to increasing drug concentrations for 4 days. Viability was assessed using CyTOX green reagent. Viability of cells with low CD271 expression was not reduced by vincristine, whereas cells with higher CD271 were sensitive to the agent **(Fig. 7E)**. We conclude that EW8 cells with low CD271 expression have altered sensitivity to vincristine.

## DISCUSSION

Knockdown of *EWSR1-FLI1* in EW8 cells revealed expression changes associated with specific subgroups of cells with different functional characteristics. Individual cells from three functionally distinct groups revealed heterogeneity in cellular responses to *EWSR1-FLI1* down-regulation and time.

We then interrogated exponentially proliferating cell EW8 cell populations and EWS xenograft tissues to determine whether cells with similar gene profiles could be identified. Our computational pipeline revealed that changes in the average expression of genes over time can dissect discriminative behaviors contributing to diverse responses. After *EWSR1-FLI1* down-regulation cells in population Dormant 2 (D2, 120h) displayed two different responses. In cluster C0, stem cell genes were upregulated, whereas they were suppressed in the C2 subpopulation **(Supplementary Fig. S11)**. Survival genes and autophagy regulatory genes were progressively induced from clusters C3 to C2, indicating an important regulatory role in maintaining this dormant state.

Following the stress induced by *EWSR1-FLI1* knockdown, the C0 cluster became the dominant population at 120h with induced stem-like gene expression and expression of inflammatory genes (*CXCL10, PTPN22, GCH1, HPSE, CCL26, PTPN6, CXCR4, AHR, PSTPIP1, BCL11B, TNF*). C0 cells showed induction of neural stem cell features defined by the expression of Cadherin EGF LAG seven-pass G-type receptor 2 (*CELSR2*) and neuron projection genes (*NPY, NEURL1, GABRA2, SYN1, PNOC, ITPKA, PCSK2, P2RY1, SNCA, CORO1A, CXCR4, BCL11B, EPB41L3, PCP4, ALK, DTNA, RND2*). These genes were associated with stem cell associated genes (*SOX2, MYC, CCND1, ALDH2*) defining the stem-like cluster C0. In mice, *CELSR2* expression is only required for the development of the forebrain at embryonic day 13.5 (E13.5) to E14.5 (49). Additional studies on mammalian cells, report a role of *CELSR2* with *PRICKEL* and Frizzled (FZD) proteins in the regulation of planer cell polarity (PCP) in epithelial sheets (50). Moreover, the C0 state showed a significantly higher level of *PRICKLE3* compared to the other 3 identified cellular states (C1, C2, C3). Interrogation of control EW8 cell populations identified ∼4% of cells with the C0 expression profile. Further rare cells were detected positive for CELSR2 by CyTOF or IHC in EWS xenografts. Functionally, cells with high expression of CD40 and CELSR2 had greater capacity to form primary and secondary spheroids in culture, characteristics of ‘stem-like’ cells, suggesting that cells in D2 with stem cell features (C0) play a role in the recovery of proliferation following *EWSR1-FLI1* knockdown.

In contrast to C0 cells, C2 cells displayed different induction strategies to remain in a proliferatively quiescent state. At 120h after *EWSR1-FLI1* knockdown, C2 cells represented 23% of the population with distinct expression patterns for genes regulating autophagy and ER, characterized by GBA, *CD63, GABARAPL2, SEC61B, RTN4, CLIMP-63* expression. Consistent with the recent single cell analysis (43) C2 and C3 cells showed lower *EWSR1-FLI1* levels and increase levels of actin-binding genes and cytoskeleton assembly and actin-based motility proteins *(MYL6, MYL12a, ACTN4, CFL1, GSN, VCL, PKP1, ITGA1, ITGA4)*. C2 cells had increased expression of genes associated with EMT, and had greater migration characteristics. Of interest, cells with the C2 profile were extremely rare in the control EW8 cell population as only one cell with this expression profile was identified in a heterogeneous (siControl, 0h) population (∼0.5%). These findings suggest that *EWSR1-FLI1* fluctuation induces a stress response and cells with the ability to suppress those signals can transition to a C0 (stem-like) state. C2 cells without the ability to suppress the ER and autophagy system remained in a proliferatively quiescent state, and hence would be anticipated to be resistant to proliferation-dependent killing by cytotoxic drugs.

In the D2 population, cells with C0 profiles (53%) or C2 profiles (24%) predominate. However, the exact cues that determine cell fate remain to be identified. The ability of C0 cells to influence other cells via paracrine signaling through secretion of *VEGFA, GDF6, BMPER, TNF, IGF1, CXCR4, CXCL10* or to be modulated through receiving signals via receptors *(FGFR4, TNFSF13B)* could be an efficient survival strategy. This idea is consistent with observations made on the relationship between cell proliferation and cell density. It indicates variability in the secretion and sensing of signals by single cells and how they are tightly regulated by inflammatory cytokines. Thus, individual cells probably display variable control on the expression of cytokines to determine cell fate under stress induced following *EWSR1-FLI1* down regulation, although the experimental conditions used here do not address this.

Using microfluidics, we were able to track gene expression changes in individual cells to identify functionally important, and rare characteristics. Consistent with the RNA-Seq findings we were able to confirm cells with stem-like (C0) expression in EWS xenografts, and the low frequency of a dormant state (C2) in EWS xenograft tissue with CyTOF.

In this study, expression of NGFR (CD271) was positively correlated with EWSR1-FLI1 expression, hence in contrast to a previous study in which EWSR1-FLI1 was shown to repress CD271 through up regulation of EZH2 (48). Down regulation of EWSR1-FLI1 resulted in decreased CD271 transcripts and protein levels in EW8 cells, and arrest of cells in G1/G0 by palbociclib also down regulated surface CD271. Importantly, a subpopulation of cells (CD271-Low) was resistant to vincristine, an agent used extensively in treatment of EWS patients. Hence, these findings may be of clinical importance to predict responses from diverse groups of EWS cells with different proliferative and survival abilities after treatment.

## Ethics Statement

Mice were maintained according to practices prescribed by the NIH at UTHSCSA Animal Facility accredited by the American Association for Accreditation of Laboratory Animal Care. All animal studies were conducted following approval from the UTHSCSA Animal Care and Use Committee (protocol 1515x)

## METHODS

### Cell Culture

Human Ewing sarcoma cells (EW8, ES4, and ES1) were established in this laboratory, TC71 was from ATCC. Cells were grown in antibiotic-free RPMI1640 medium supplemented with 20% FBS (Sigma). Cells were maintained at 37 C with 5% CO_2_ and 90% humidity. For single-cell RNAseq, 5×10^5^ cells were seeded in each 100-mm dish and cultured overnight. After 24h, cells were treated with 50nM human On-TargetPlus si-EWSR1 (Cat#J-005119-11-0050; Dharmacon, General Electric) or 50nM siControl (Accel Control Pool, Non-Targeting, Cat#D-0019; Dharmacon, General Electric) in Dharmafect1 (Cat#T-2001-03; Dharmacon). The target sequence for EWSR1 was “GAGUAGCUAUGGUCAACAA”.

### EWSR1 Rescue

To over express mutant EWSR1 with two silent mutations that protected from knockdown by siEWSR1, we cloned EWSR1+mCherry into the tetracycline inducible plasmid pLIX_403 vector (pLIX_403 was a gift from David Root, Addgene plasmid # 41395). This clone was confirmed by sequencing and named PLIX-EWSR1-2 mut. The lenti-X packaging single shot system was used to package virus. Virus was grown in human embryonic kidney HEK293T, and media-containing virus was passed through a 0.45 μM filter and stored at −80 C. EW8 cells were infected with virus, and selected in puromycin. Cells were treated with 2.5 μg/ml doxycycline to express EWSR1.

### C1 microfluidic-based single-cell RNA-Seq and data analysis

Single EW8 cells were isolated with the C1 microfluidic system. Three 96 well chips were used for each condition (Control, Dormant 1, Dormant 2) and a C1 SMART-Sq V4 protocol was used for the reaction mixes and thermo-cycling. The Fluidigm modification of the Nextera XT kit was used to prepare libraries for 864 cells (Fluidigm 100-7168). Ethidium homodimer-1 (in LIVE/DEAD Viability/Cytotoxicity Kit, for mammalian cells, Life Technologies, PN L-3224) was used to distinguish dead cells before library preparation. ERCC RNA Spike-In Mix 1 (Cat#4456740;Life Technologies) was used to normalize raw sequencing data and reads aligned to UCSC hg19 were converted to transcripts per million (TPM) to quantitate gene expression level. High quality cells were selected based on a threshold for the number of genes detected in each individual cell (a minimum of 2000 genes per cell). Data were log-transformed [log (TPM+1)] for data analyses and visualization. Live single-cell libraries were sequenced with HighSEQ 3000 (Illumina) and three technically matched culture replicates were prepared for all three-time points (0h, 48h, 120h). Each library was sequenced to an average depth of 2×10^6^ read pairs. A range of 7000-9000 genes was detected in each cell.

### Single-cell profiling and cluster discovery

To identify novel sub-clusters of cells with discriminative markers, transcriptionally perturbed cells with siEWSR1 were pooled and feature selection method (44) was applied to the data generated from two time points (48 and 120h). Novel surface markers were identified from each key cluster by KEGG pathway enrichment analysis was performed using (http://www.genome.jp/kegg/pathway.html) to identify the top biological processes in the gene sets based on the KEGG database. The pathways with p-values <0.001 and fold change >2 was used to determine significant cellular processes enriched in each sub-cluster.

### Gene expression variation model parameter estimation

We assumed gene expression possesses a Gaussian mixture model with up to 2 modes: one is from cells that the gene is not expressed (log2 (TPM+1) ∼ 0) while the other is from cells that the gene is activated (log2 (TPM+1) > 1). We then fit the model by maximum likelihood, using the Expectation-Maximization (EM) algorithm, implemented by fitgmdist function of MATLAB to estimate 3 nominal parameters, *μ*, *σ*^2^, (for mean, variance) and *α* for each gene in each condition (control, dormant 1 and dormant 2). Log2 (TPM+1) was used for expression levels. We define *a* value as the proportion of cells where transcript is expressed (or detected at level log2 (TPM+1) >1).

### Gene activation and induction score

We used logistic regression to establish a relationship between the detection rate and expression level. Specifically, for each cell, we define following function (58),

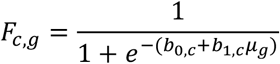

Where *b*_0,*c*_ and *b*_1,*c*_ are the fitted parameters for the logistic regression for cell *c*, and *μ_g_* is the expression level for gene *g* in cell *c*.

#### Induction Score

We then calculated average expression levels for all genes in the control population as *μ_control,g_*. For a given cell, we transferred expression levels into “induction value” by dividing by the average expression of the gene in the control population. To reduce technical variability, we weighted each gene by the activation function obtained from logistic regression given before (41), or the final induction score 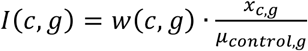 for a given cell *c* and gene *g*, where

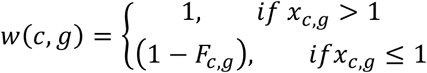

and *x_c,g_* is the gene expression level in log2(TPM+1).

#### Functional module induction score

A functional module induction score *M*(*c, S*) of a cell was then calculated as the weighted mean of “induction values” over a set of gene *S* (functional gene set) within the cell, or

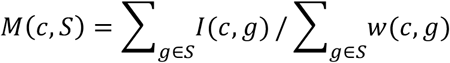

In this study, 3 functional gene sets were considered, cell-cycle, stemness, and dormancy (Supplemental Table S1).

### Differential Induction score, heat map, and functional enrichment

Student *t* tests were performed to identify dormant populations (siControl, Dormant 1 and Dormant 2), or 4 class-labeled clusters (C0, C1, C2, and C3). Significant genes with differential induction scores were selected based on: 1) Benjamini-Hochberg multiple-test adjusted p-value < 0.05; 2) Induction score > 0.2; and 3) at least 2-fold-change in induction score. Genes with differential induction scores were collected to generate a heat map (highest induction score in red-color, and low induction score in white). Genes and cells were sorted based on their average induction score within each group and across cells, respectively. In addition, genes were grouped to each group’s class-label if their mean induction scores were higher in the group than other groups. Functional enrichment was performed using *enrichr* website.

### Western blotting

Cells were lysed in RIPA buffer and run in 4–12% Mini-NuPage Bis-Tris precast protein gels (Thermofisher). The primary antibodies used were rabbit anti-EWSR1 (no.11910S; Cell Signaling), rabbit anti–FLI1 7.3 (SC-53826,LOT#0213; Santa Cruz) and rabbit-anti-FLI1 (ab15289;Abcam), rabbit anti–GAPDH (D16H11;Cell Signaling).The secondary antibody used was anti-rabbit IgG HRP-linked antibody (7074S; Cell Signaling). Cell treatment, sample collection, and Western blotting were repeated at least three times, and the representative blots are shown in the figures.

### Flow cytometry studies and cell sorting

PE mouse anti-human CD271 (Cat#557196), PE mouse anti-Human CD40 (Cat#555589), PE mouse anti-Human CD63 (Cat#556020) were purchased from BD Biosciences for FACS analyses. The Quantibrite PE beads (Cat#340495; BD Biosciences) were used to profile cells with Low-Medium-High expression levels. Cells were sorted into 4 sub-clusters based on their expression profiles for markers CD271 (PE Mouse Anti-Human CD271, Cat#5571196; BD Pharmingen), CD63 (PEMouse Anti-Human CD63, Cat#556020; Bd Pharmingen), CD40 (PE Mouse Anti-Human CD40, Cat# 555589; BD Pharmingen).

### Immunofluorescent imaging

Cells were grown in chamber slides (MatTek; Thermo Fisher) and treated with siEWSR1 for 48h and 120h. Cells were fixed in 100% methanol and blocked with 5% goat serum. Primary antibodies were Alexa Fluor 647 anti-human CD63 (Cat#561983), LC3A/B (D3U4C) rabbit mAB Alexa-555 (131 Cell Signaling), anti-FAM134B (HPA012077; ATLAS), anti-CELSR2-N-terminal (ab189045). The secondary antibody Alexa Fluor 488 (ab150077; Abcam) against FAM134B, SEC61B (sc-393633; Santa Cruz) was used. Cells were stained and mounted with ProLong Gold Antifade mountant with DAPI (P36931; Thermofisher Scientific). Images were taken using a Flouview FV3000 confocal laser-scanning microscope (Olympus). Signal analysis was performed with FIJI software (GPL v2;ImageJ).

### CyTOF and Immunostaining

In vitro assessments of protein expression variations in response to EWSR1-FLI1 down-regulation were studied using CyTOF (Helios 2; Fluidigm Inc.) and acquired data were analyzed using FlowJo software v10.2 (Tree Star Inc.) and Cytobank (Cytobank Inc.). EW8, ES4, ES1 cells treated with siEWSR1 were selected after 48h and 120h knockdown. Two vials of each line were obtained for the no staining control and functional control (si-Control, 0hr). All cells were treated with 5 μM Cell ID Cisplatin (Fluidigm; Cat# 201064) in polypropylene round-bottom tubes, 5 mL capacity (Becton Dickinson 352235) for 5 min. Staining protocol for antigens located in the nucleus and cell surface markers was applied (Fluidigm) to 3×10 ^6^ cells in 100 μl staining volume. The primary antibodies used were human Cyclin B1-153Eu (Fluidigm; 3153009A), Cyclin D-141Pr (Cell Signaling;2978BF), CD271-149Sm (Fluidigm;3149017B), CD40-142Nd (Fluidigm; 3142010B), CD79a-158Gd (Cell Signaling;13333BF), CD49b-161Dy (Fluidigm; 3161012B), CD49e-160Gd (Fluidigm; 3160015B), CD63-150Nd (Fluidigm; 3150021B), CDKN2A-155Gd (Abcam;Ab186932), CELSR2-165Ho (Abcam; Ab189045), FGFR1-159Tb (Cell Signaling;9740BF), MKI67-147Sm (Cell Signaling; 9449BF), MYC-176Yb (Fluidigm;3176012B), TGFBR2-170Er (Abcam;Ab 78419),TGFB1-163Dy (Fluidigm;3163010B), SOX2-169Tm (Cell Signaling;3579BF). To prevent nonspecific background signal, cells were blocked with Fc protein that blocks nonspecific binding of Fc receptor expressing cells with antibody (Cat#564220; BD Pharmingen) for 30 min before staining with 1.5 μL of each antibody over night at 4 ° C. Cells were labeled with 125 nM Intercalation solution (Fluidigm; Cat#201192B) for 1h at the room temperature. Cells were washed after each step for 5 min and centrifuged at 350Xg before cell fixation for 5 min and centrifuged at 1000Xg after cell fixation. Pelleted cells were re-suspended in Maxpar water (Fluidigm; Cat# 201069) for data acquisition with the mass cytometer Helios2 in the Flow Cytometry and Cellular Imaging Core Facility at the University of Texas MD Anderson Cancer Center.

### Cell proliferation study and viability assays

Cells plated at identical densities (5000 cells/well) were incubated in 96-well plates (Cat#3596; Corning). Cell proliferation was assessed by measuring confluence (%) over 96h with the IncuCyte system (Essen Bioscience Inc.) Images were taken every 4h from 4 different areas of each well. To distinguish live from dead cells the Cytox green reagent was added to each well at the end point to yield a final concentration of 250nM. Images were taken from 3 technical replicate and studies were repeated at least 3 times.

### Functional studies for the stem-like cluster, flow cytometry staining and cell sorting

Self-renewal studies were conducted on FACs sorted cells based on the CD40 signal. Cells isolated based on their CD40 expression level were plated at identical densities (50 cells /well) and incubated in 96-well plates (Ultra Low Cluster round bottom, ultra-low attachment polystyrene, Cat#7007, Corning). Cells were grown in knockout DMEM/F-12 (Gibco;Life technologies)) supplemented with 1% penicillin-streptomycin (Thermo Fisher Scientific) and with EGF, FGF growth factors, and StemPro Neural supplement (Cat#A1050801;Thermo Fisher Scientific) for 7 days. At day 7, images were taken with the Celigo S imager (Nexcelom Bioscience) to assess the ability of each population to form spheres with the Celigo sphere-formation tool.

### Endoplasmic reticulum (ER) area measurements

ER area measurements were conducted using ImageJ as follows. Background threshold was manually defined and set for all images. Borders of each cell were drawn and the ER area was then calculated and presented as a fraction of the total cell area.

## Contribution

Conception and design: RKH, PJH

Development of methodology: RKH, PJH

Acquisition of single-cell RNA seq and CyTOF data: RKH

Pipeline design and cell state assignment: RKH

Functional analyses, immunofluorescence and cellular localization: RKH

Established PLIX-EWSR1 cell line (EW8): RKH

Interpretation and conclusions: RKH, PJH

Statistical analysis and result discussion: RKH, PJH, YC, ERL, KK, MI.

Writing, review and/or revision of the manuscript: RKH, ERL, YC, KK, MI, TH, PJH.

Administrative, technical, or material support: PJH

Study supervision: PJH, YC, MI.

## Acknowledgments

This study was supported by USPHS award PO1CA169368 (PJH) form the National Cancer Institute. We thank the core staff and investigators involved in data collection. Single-cells were isolated in the UTSA Genomics Core, which is supported by UTSA, NIH grant G12MD007591, and NSF grant DBI-1337513.

Single-cell RNA-Seq data were generated in the Genome Sequencing Facility at GCCRI, which is supported by UT Health San Antonio, NIH-NCI P30 CA054174 (Mays Cancer Center at UT Health San Antonio), NIH Shared Instrument grant 1S10OD021805-01 (S10 grant), and CPRIT Core Facility Award (RP160732).

CyTOF data were generated in the MD Anderson Cancer Center Flow Cytometry and Cellular Imaging Core Facility, which are partially funded by NCI Cancer Center Support Grant P30CA16672 and directed by Dr. Jared Burks.

Flow data were generated in the Flow Cytometry Shared Resource Facility, which is supported by UTHSCSA, NIH-NCI P30 CA054174 (CTRC at UTHSCSA) and UL1 TR001120 (CTSA grant) directed by Mrs. Karla Gorena.

---

**Supplemental Figure S11.**
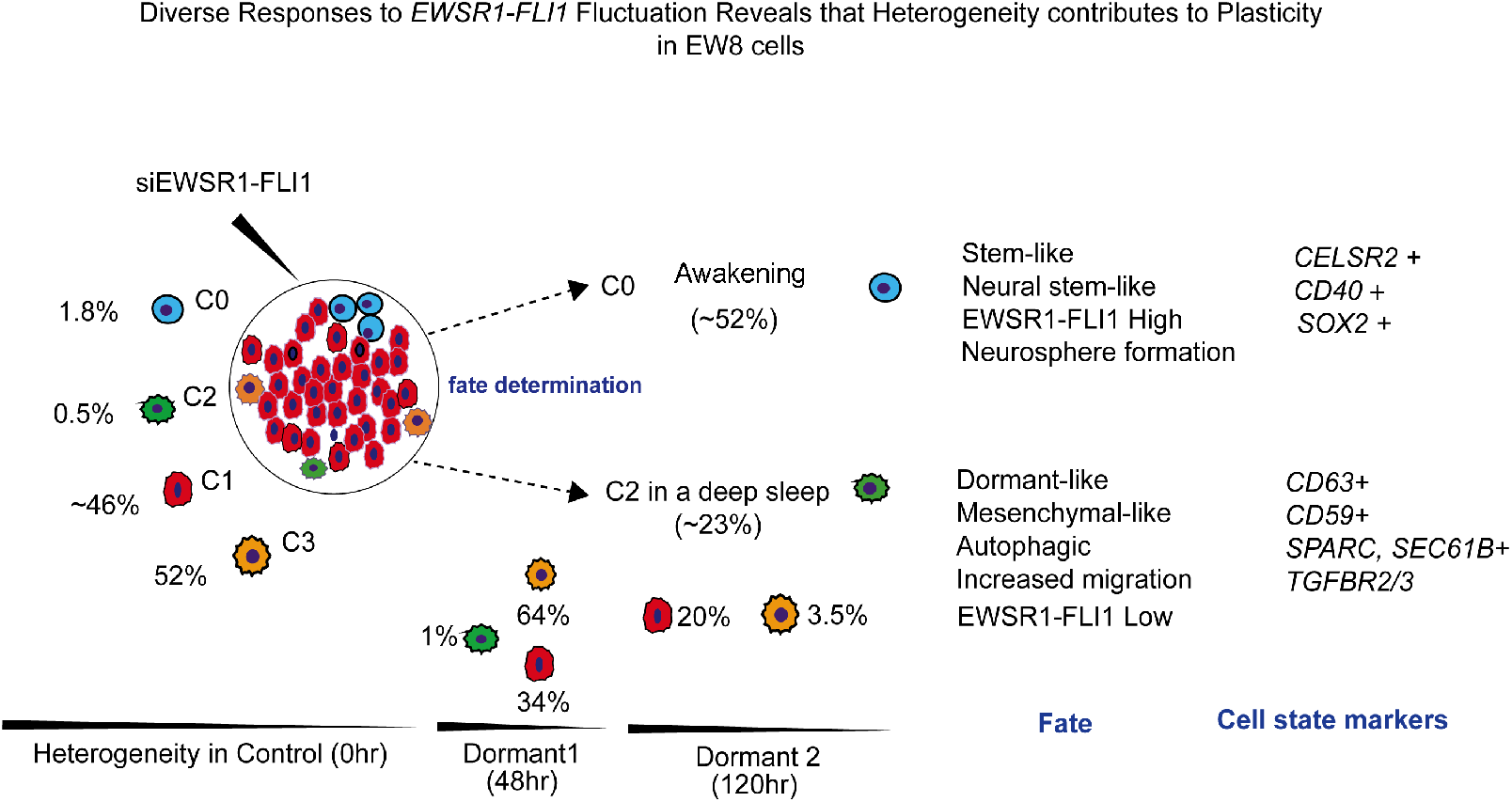
An overview of the frequency of 4 newly identified cell states in EW8 cells following EWSR1-FLI1 perturbation. In this schematic figure we show frequency changes of the 4 subclusters (C0, C1, C2, C3) in three different time points (0, 48, 120h) following the identical perturbation of EWSR1-FLI1 suppression (48h and 120h).

